# Few-Shot Viral Variant Detection via Bayesian Active Learning and Biophysics

**DOI:** 10.1101/2025.03.12.642881

**Authors:** Marian Huot, Dianzhuo Wang, Jiacheng Liu, Eugene Shakhnovich

**Affiliations:** Department of Chemistry and Chemical Biology, Harvard University, Cambridge, MA; Laboratory of Physics of the Ecole Normale Supérieure, CNRS UMR 8023 and PSL Research, Sorbonne Université; John A. Paulson School of Engineering and Applied Sciences, Harvard University, Cambridge, MA; University of Washington, Seattle, WA

**Keywords:** Pandemic Prevention, Protein Evolution, Active Learning, Antibody Escape, Protein Language Models

## Abstract

The early detection of high-fitness viral variants is critical for pandemic response, yet limited experimental resources at the onset of variant emergence hinder effective identification. To address this, we introduce an active learning framework that integrates protein language model ESM3, Gaussian process with uncertainty estimation, and a biophysical model to predict the fitness of novel variants in a few-shot learning setting. By benchmarking on past SARS-CoV-2 data, we demonstrate that our methods accelerates the identification of high-fitness variants by up to fivefold compared to random sampling while requiring experimental characterization of fewer than 1% of possible variants. We also demonstrate that our framework benchmarked on deep mutational scans effectively identifies sites that are frequently mutated during natural viral evolution with a predictive advantage of up to two years compared to baseline strategies, particularly those enabling antibody escape while preserving ACE2 binding. Through systematic analysis of different acquisition strategies, we show that incorporating uncertainty in variant selection enables broader exploration of the sequence landscape, leading to the discovery of evolutionarily distant but potentially dangerous variants. Our results suggest that this framework could serve as an effective early warning system for identifying concerning SARS-CoV-2 variants and potentially emerging viruses with pandemic potential before they achieve widespread circulation.

## Introduction

The relentless ascent in the number of SARS-CoV-2 infection cases has catalyzed an unprecedented protein evolution, yielding a profusion of novel mutations. Mutations in the receptor-binding domain (RBD) of the viral spike protein are particularly consequential, as they can increase binding affinity to the ACE2 receptor Ozono et al. (2021); Barton et al. (2021), facilitating more efficient host cell entry, or reduce susceptibility to neutralization by monoclonal antibodies (mAbs) and convalescent sera Tuekprakhon et al. (2022); Nabel et al. (2022); Wang et al. (2023). These adaptations enable the emergence of lineages with higher fitness Carabelli et al. (2023), which are better equipped to spread within populations or evade immune responses, thereby posing significant challenges to public health interventions and vaccine efficacy. As we navigate the aftermath of this crisis, it is imperative that we strengthen our preparedness for future pandemic threats.

Supervised machine learning has emerged as a valuable tool for predicting viral fitness (infectivity) and identifying high-risk viral mutations. These models leverage fitness labels collected from sequencing databases such as GISAID (Elbe and Buckland-Merrett, 2017a) or integrate laboratory measurements with epidemiological data. For example, Obermeyer et al. Obermeyer et al. (2022) introduced a non-epistatic approach that infers fitness from GISAID, allows estimation of fitness for combination of observed mutations. Ito et al. Ito et al. (2024) and Maher et al. Maher et al. (2022) integrate laboratory measurements of binding affinities with epidemiological data to train machine learning models that forecast the fitness of the SARS-CoV-2 variants and identify variants susceptible to widespread transmission. However, these models were rare in the early stages of the pandemic due to the limited availability of labeled fitness data. Fitness labels are difficult to obtain unless the virus has been propagating in the population for a sufficient period.

Given these limitations, biophysical models offer a complementary approach by linking experimentally determined binding constants (*K*_*D*_) to fitness as demonstrated by these works Wang et al. (2024); Chéron et al. (2016); Rotem et al. (2018). A key advantage of biophysical modeling is its ability to significantly reduce the functional space compared to machine learning, enabling accurate predictions with minimal experimental data while maintaining as a powerful fitness predictor. The underlying intuition is that high-fitness variants are usually variants that evade antibody binding while maintaining strong cellular receptor affinity. These experimental values can be obtained through low-throughput methods such as surface plasmon resonance Zhang et al. (2024) and isothermal titration calorimetry Upadhyay et al. (2023) or high-throughput methods such as deep mutational scanning (DMS) Starr et al. (2020) or combinatorial mutagenesis (CM) Moulana et al. (2023). However, the time and cost associated with comprehensive experimental measurements can limit its availability during the early stages of a pandemic.

Active learning addresses the challenge of limited labeled data by prioritizing the most promising variants for experimental characterization and has been successfully used for discovery of high fitness proteins Hie et al. (2020) as well as chemical space exploration Khalak et al. (2022); Graff et al. (2021). Active learning framework can efficiently be combined with Gaussian Process (GP), as they excel at handling scarce labeled data while providing uncertainty estimates Hie et al. (2020); Romero et al. (2013); Gessner et al. (2024).

This paper explores the integration of protein language model (pLM), active learning and biophysical modeling to enhance early pandemic response capabilities, including potential variants of concern (pVOC) detection, and identifying sites under-pressure. We combine a biophysical model Wang et al. (2024) based with a machine learning pipeline that integrates active learning with GP decoder on embeddings acquired from ESM3. ESM3 is a state-of-the-art pLM that generates structureaware sequence embeddings Hayes et al. (2024), which the GP uses to smoothly predict how mutations affect binding affinities to cell receptors and antibodies. Then these binding affinities are piped into a pre-trained biophysical model Wang et al. (2024). This approach efficiently identifies potentially dangerous mutations by prioritizing the most informative variants based on acquisition functions.

We validated this pipeline using deep mutational scanning data, combinatorial mutagenesis experiments, and GISAID sequencing data, demonstrating that our few-shot learning approach can effectively substitute for high-throughput screening in early pandemic surveillance.

## Results

### A. Overview of VIRAL

We introduce VIRAL (Viral Identification via Rapid Active Learning), a framework that combines GP with active learning to predict the binding specificity of SARS-CoV-2 Receptor Binding Domain (RBD) variants to ACE2 and various antibodies (Figure 1a). The model operates in a structured pipeline: first, the RBD structure and mutant sequence are input into ESM3, a protein language model, to generate sequence embeddings that incorporate both sequence and structural context. These embeddings are then fed into a GP trained on a limited set of experimental dissociation constants to predict binding affinities. The predicted dissociation constants, as well as predicted uncertainties, are subsequently passed into into a biophysical model, following our previous work Wang et al. (2024), to infer the infectivity of each variant.

**Figure 1.**
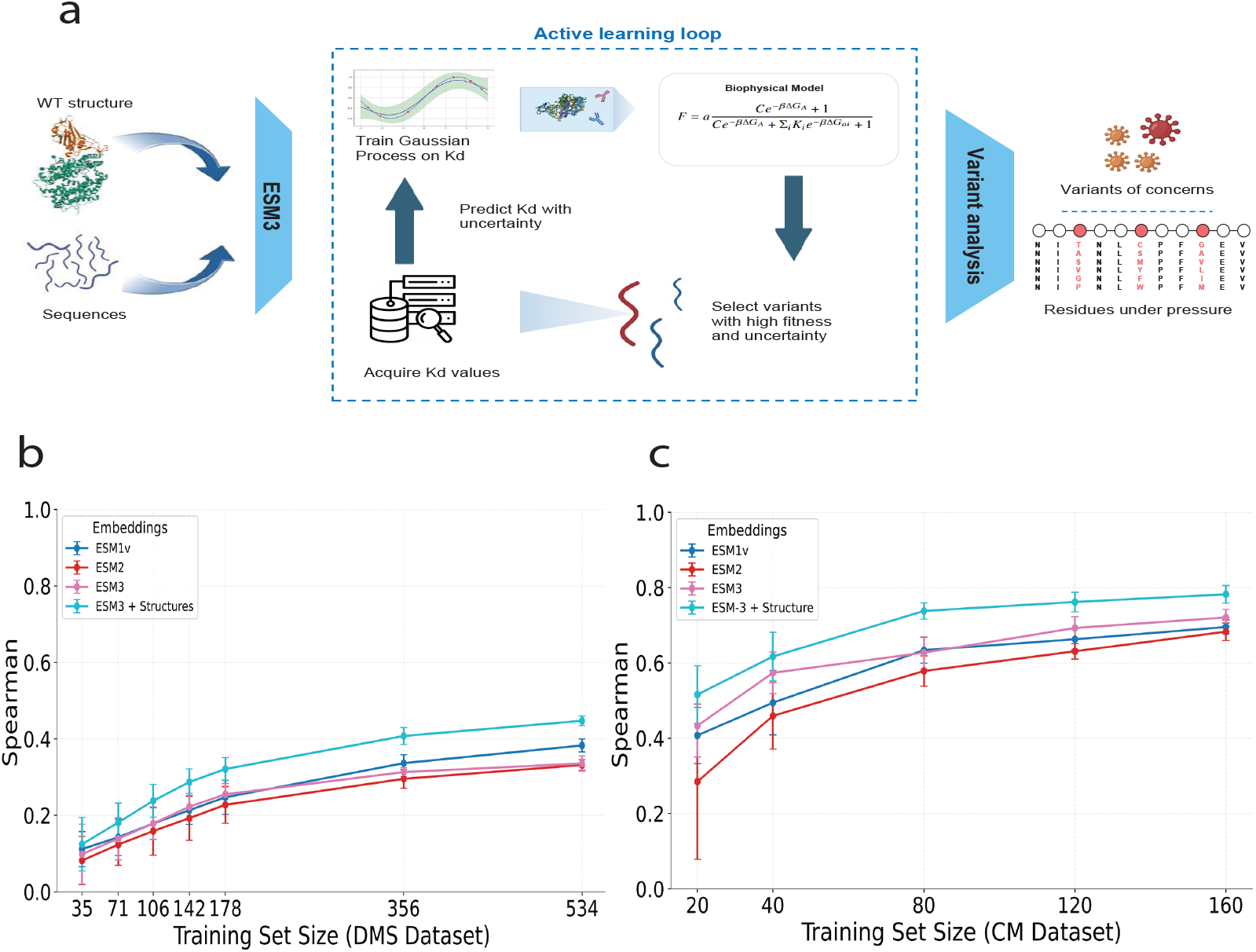
Overview of active learning framework for detecting high-fitness SARS-CoV-2 RBD variants. (a) The pipeline begins by using ESM3 and RBD structure to generate embeddings for RBD sequences. These embeddings serve as inputs to a combined Gaussian Process and biophysical model that predicts variant fitness. The framework operates in an iterative cycle where: (1) The model predicts fitness and uncertainty for untested variants, (2) Based on these predictions, the most promising variants are selected for experimental testing using various acquisition strategies (greedy, UCB, or random sampling), (3) An oracle representing experiments provides binding constants (Kd values), and (4) These new measurements are used to retrain the GP model, improving its predictive power. (b) Spearman correlation of predicted fitness for different training sized on DMS dataset. Error bars show std. Esm3_coord refers to ESM3 with wildtype structure. (c) Similar to (b) but on CM dataset.

A key strength of our approach is its few-shot capability—the ability to rank variant fitness effectively even with extremely limited labeled data points, making it particularly powerful for early variant detection during emerging outbreaks. This capability leverages the Gaussian Process’s inherent ability to learn efficiently from small *K*_*D*_ datasets by utilizing the prior knowledge encoded in its kernel function, while simultaneously providing valuable uncertainty estimations about its predictions. Seeger (2004) Additionally, our biophysical model can be trained with minimal data to provide an effective mapping from *K*_*D*_ to fitness. Wang et al. (2024)

Notably, in our benchmark studies (Figures 1b, 1c), incorporating structural information from the wildtype RBD (PDB 6XF5 Zhou et al. (2020)) to ESM3 significantly enhances predictive accuracy across both deep mutational scanning and combinatorial mutation benchmarks compared with sequence-only methods such as ESM3 sequence only, ESM2, and ESM1v. This advantage of structure-aware embeddings is consistent with Loux et al.. In particular, our ESM3 with structure achieves a spearman coefficient of 0.53 on combinatorial dataset while being trained on only 20 points (0.06% of dataset). Based on these results, all subsequent analyses presented in this work utilize the ESM3 embeddings that integrate both sequence and structural information.

The predictions from this pipeline guide an active learning strategy: variants with high predicted infectivity are selected for validation, and new measurements are used to iteratively retrain the GP, refining its accuracy over time. This iterative process enables the model to rapidly identify high-risk variants while minimizing the number of required measurements, making it a scalable approach for early warning systems in viral surveillance.

### B. Active learning identifies top variants

Our objective is to maximize the identification of potential Variants of Concern (pVOC), defined as those ranking in the top p=10% across the mutational landscape. To rigorously evaluate our approach, we start with a retrospective study using existing Combinatorial Mutagenesis (CM) datasets containing experimentally measured binding constants (see Methods). This retrospective analysis is crucial as it provides the only systematic way to benchmark our model’s performance: by simulating the sequential selection of variants from a completely characterized fitness landscape, we can precisely quantify how efficiently our framework identifies high-fitness variants compared to alternative strategies. Such comprehensive validation would be impossible in a prospective setting, where the fitness of untested variants remains unknown.

Our main analysis focuses on two acquisition strategies: a greedy strategy that selects variants based solely on predicted fitness, and an Upper Confidence Bound (UCB) approach that combines predicted fitness with model uncertainty (see Method). While the greedy strategy excels at exploiting regions of known high fitness, UCB balances exploitation with exploration of uncertain regions in the sequence landscape, potentially uncovering novel fitness peaks. Random sampling constitutes the baseline, mimicking what we could achieve using brute force searching—screening every RBD in a library indiscriminately. We define the enrichment factor (EF) as the ratio of the percentage of top variants found by the model-guided search to the percentage of top variants found by a random search. An EF greater than 1 indicates superior performance compared to brute-force screening.

We use an initial training set of variants with a maximum number of 2 mutations, to model the first observed variants in a pandemic. Given that the most concerning variants, such as BA.1, can accumulate up to 15 mutations in the Receptor Binding Domain (RBD), an effective early warning system must efficiently evaluate variants with increasingly complex mutation combinations to assess their potential for complete antibody escape. This translates into UCB acquisition metric achieving a final EF of 5 after 10 rounds of acquisition corresponding to a total of 120 points and 0.4% of the dataset (Figure 2b), while the greedy exploration struggles to beat random baseline: lack of exploration translates into lower capacity to identify dangerous variants (Figure 2a), quantified by Area Under the Receiver Operating Characteristic Curve (AUC).

**Figure 2.**
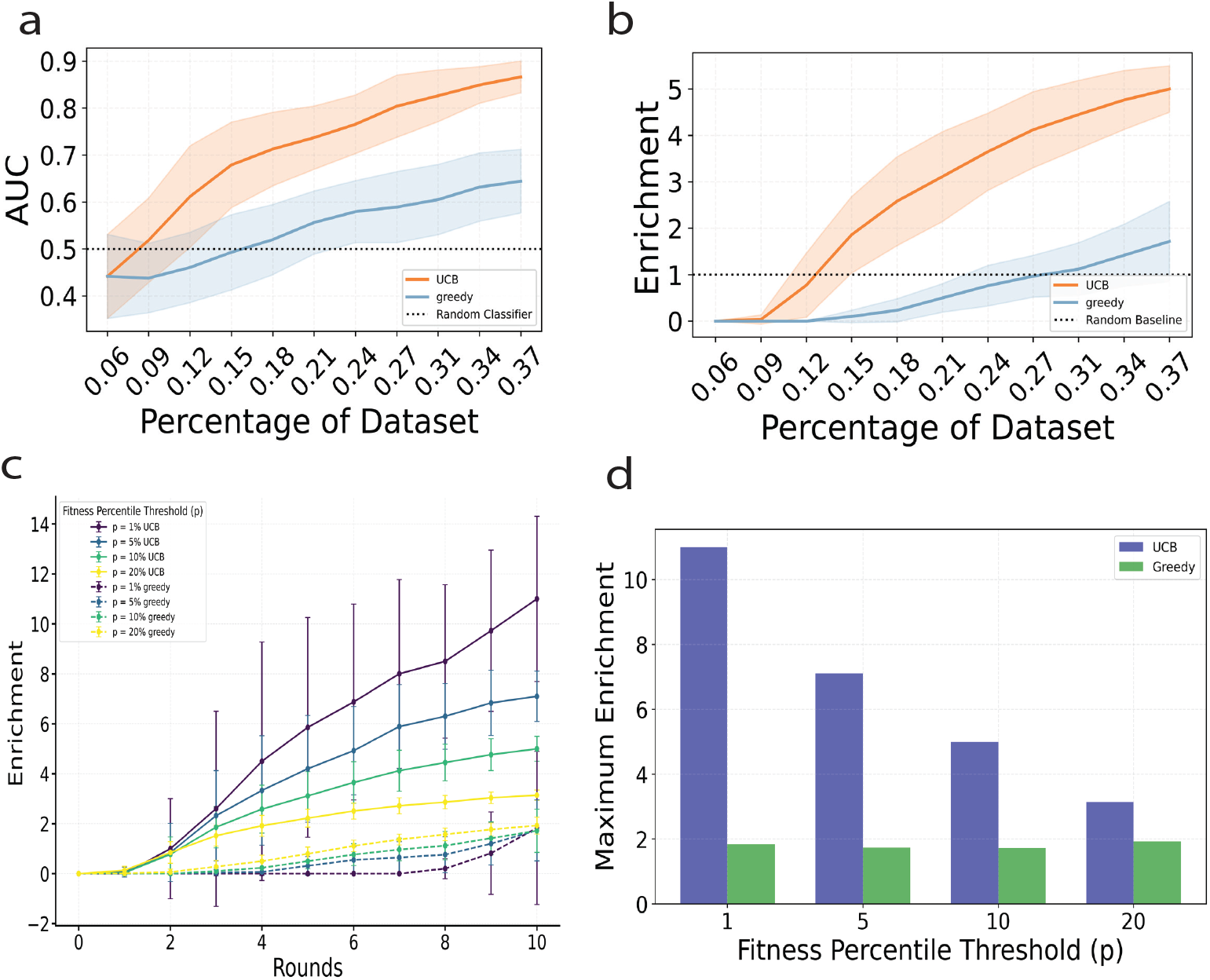
Active learning performance on combinatorial dataset. (a) Area Under the Curve (AUC), representing the model’s ability to identify top fitness variants, is shown for different strategies (UCB and greedy). An AUC above 0.5 indicates effective identification of high-fitness variants. Each round corresponds to acquiring a new batch of variants, improving the predictor. Shaded regions represent standard deviations across runs. (b) Enrichment in top variants across acquisition runs for each strategy. Values above 1 represent improvement over a random acquisition. (c) Enrichment across acquisitions runs for different fitness thresholds *p* defining top variants. (d) Maximum enrichment obtained during active learning for different fitness thresholds *p* defining top variants.

The performance of the model improves when we increase the stringency of our definition for dangerous variants by decreasing the threshold *p* (Figures 2c and 2d). For instance, when defining dangerous variants as those in the top 1% of fitness scores rather than the top 10%, the model achieves a maximum enrichment factor of 11, demonstrating particularly strong performance in identifying the most concerning variants.

### C. Exploration of dataset

The contrasting enrichment factors between UCB and greedy strategies reflect their fundamentally different exploration behaviors. Figure 3a illustrates this distinction through the exploration variance in the ESM3 embedding space, where UCB demonstrates substantially higher variance of the embeddings for acquired points compared to the greedy approach (see SI Figure S2 for impact of uncertainty weight in UCB exploration). This increased variance signifies that UCB conducts a more thorough and diverse exploration of the sequence landscape, systematically sampling from a broader range of potential variants rather than concentrating on a limited region of the sequence space. This broader exploration is visually demonstrated in the UMAP visualization (Figure 3b), where UCB successfully identifies and samples from multiple distinct clusters of high-fitness variants, while the greedy strategy remains confined to a more limited region of the sequence space. This observation highlights UCB’s ability to balance exploitation of known high-fitness regions with exploration of potentially promising but unexplored sequence clusters, in contrast to the greedy strategy’s more localized search pattern.

**Figure 3.**
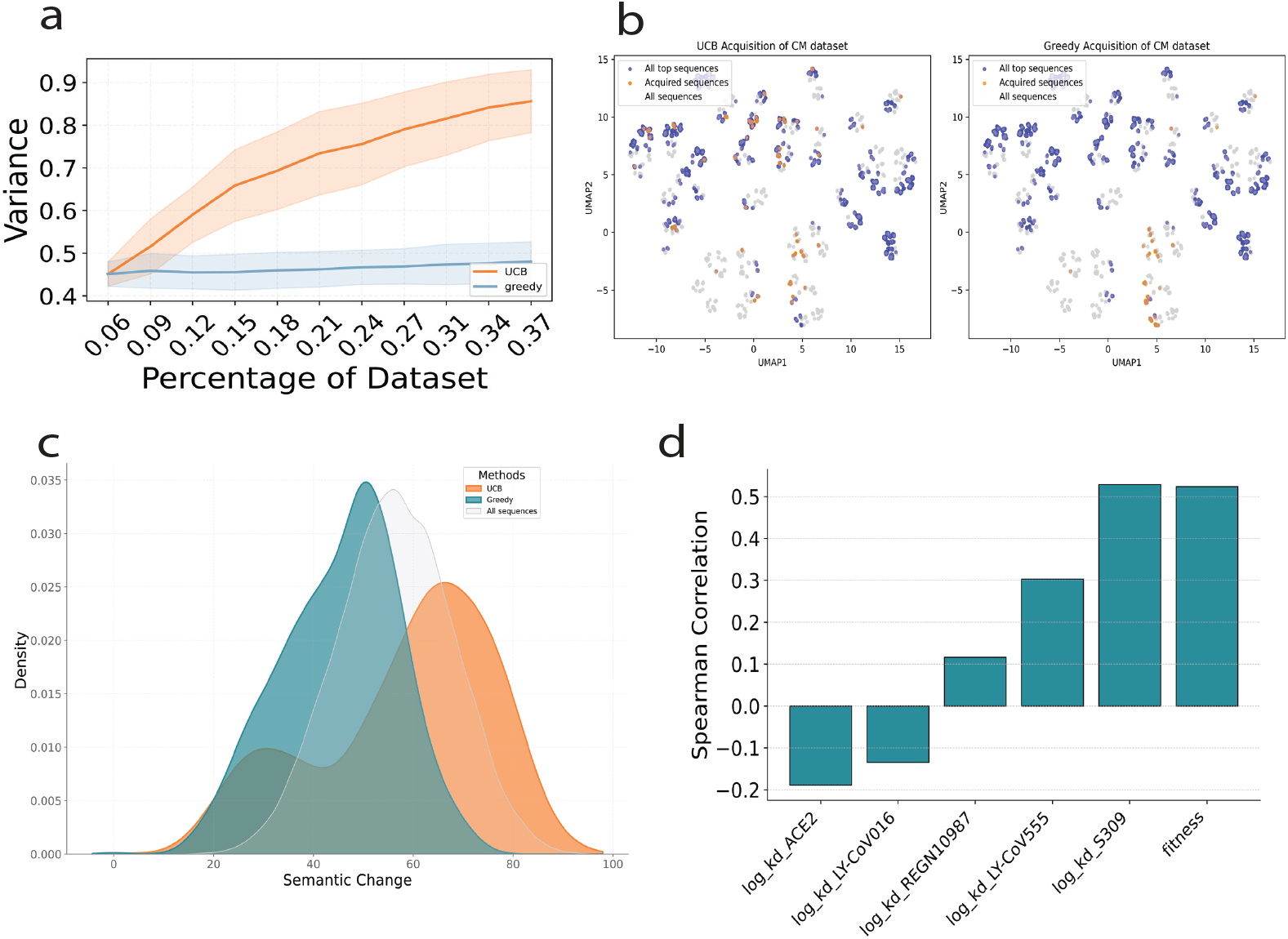
Exploration of combinatorial landscape (a) Comparison of embedding variance of points acquired across rounds between UCB (orange) and greedy (blue) strategies. (b) UMAP visualization of sequence space comparing acquired variants (orange) against all top sequences (blue) and background sequences (gray) using UCB (left) and greedy (right) acquisition strategies. (c) Distribution of semantic change of acquired RBD variants using different strategies. (d) Correlation of ACE2/antibody binding and fitness with semantic change.

In Figure 3c, the density plot reveals the semantic change distribution of sampled variants. Semantic change is defined as the Euclidean distance between wildtype and mutant (see methods). We observe that UCB’s broader sampling extends into regions shifted towards higher semantic change. This broader sampling is particularly important, as shown in Figure 3d, where higher semantic change positively correlates with enhanced immune escape and increased viral fitness. The ability to explore sequences with greater semantic divergence from wild-type is crucial, as these more distant regions of sequence space often harbor novel beneficial mutations that could drive the emergence of escape variants.

### D. VIRAL identifies highly mutable sites

Our framework effectively identifies sites that emerged as mutation hotspots during natural viral evolution, particularly those linked to antibody escape. A site is classified as “highly mutable” if, over the entire course of pandemic, it has exhibited more than *threshold* = 9 out of 20 possible amino acid substitutions. (See SI Figure S3 for model performances with different thresholds)

After running VIRAL on the DMS dataset with different initial training sets (maximal enrichment of 3.1 and AUC of 0.81, see SI Figure S1), we found a strong correlation between sites frequently sampled by our algorithm and those that became highly mutable in natural evolution. Across 10 independent runs with different initial training sets, the AUC values ranged from 0.62 to 0.78. When acquisition scores were averaged across multiple runs to reduce sampling noise, we obtained a robust AUC of 0.76 (Figure 4a). These results remain consistent across different thresholds used to define “highly mutable” sites in natural evolution (SI Figure S3).

**Figure 4.**
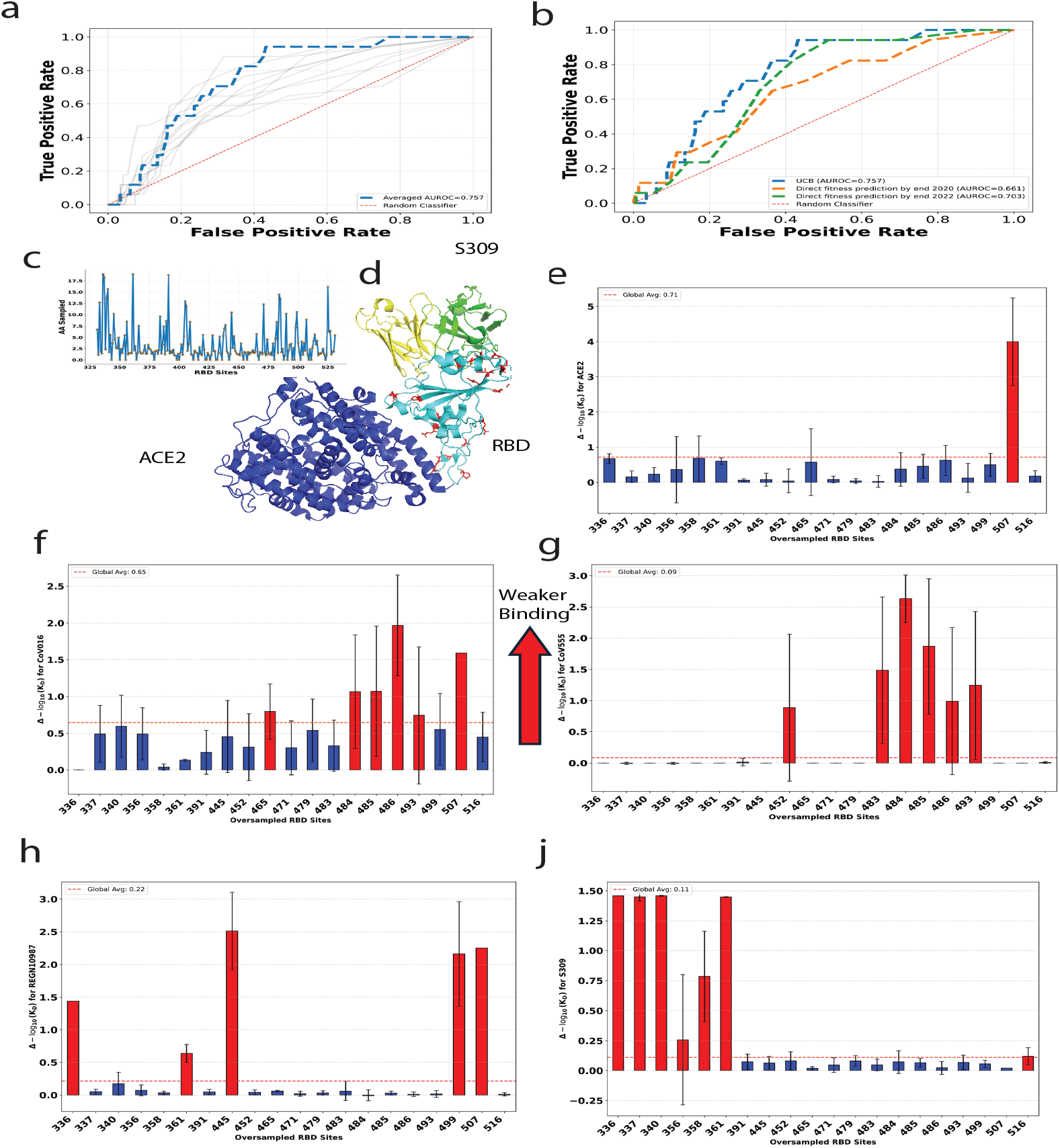
Active learning identifies residues under selection pressure. (a) ROC curves illustrating the performance of VIRAL in identifying frequently mutated sites in the GISAID database. Individual experimental runs are shown as grey lines. Blue curve is obtained when averaging the acquisition scores across multiple runs to reduce sampling noise. (b) Comparative ROC analysis of VIRAL benchmarked against one-batch acquisition using single-mutation variants observed by the end of 2020 (orange) or 2022 (green). (c) Number of sampled amino acid per site, plotted against the RBD sequence. (d) Structural representation of the RBD (light blue) in complex with ACE2 (dark blue) and the S309 antibody (green and yellow, representing the heavy and light chains, respectively; PDB: 8FXC). Red highlights indicate the top 20 sampled sites by VIRAL. (e-j) The KD distribution for oversampled sites identified by VIRAL is shown for ACE2, LY-Cov016, LY-CoV555, REGN10987, and S309. Higher values indicate binding loss. Red-highlighted sites cause escape and exhibit a greater average mutational binding loss than the protein-wide average.

We bench-marked our methodology against alternative approaches to predict mutation-prone sites in Figure 4b. Our integrated approach significantly outperforms baseline predictions derived from variants observed before end of 2020, while performing slightly better than to predictions based on variants until end of 2022. This competitive performance is particularly noteworthy because baseline predictors have an inherent advantage—they are trained directly on pandemic-era mutations that successfully emerged in the population, while our method identifies relevant sites without this prior knowledge. Precisely, with initial training set of only one single variant per site, our pipeline enhances predictive capability by up to two years in identifying sites under pressure.

Figures 4c and 4d provide sequence and structural representation on the sites prioritized by our framework. The sampling frequency across the RBD sequence is visualized in Figure 4c, while Figure 4d maps the top 20 most frequently sampled sites onto the structure of the RBD-ACE2-S309 complex (PDB: 8FXCAddetia et al. (2023)). The spatial distribution of these prioritized sites reveals a immunological pattern: they cluster in critical regions, the ACE2 binding interface (which overlaps with class 1 and 2 antibody epitopes) and regions recognized by class 3 and 4 antibodies.

Figures 4e–j further analyze the functional impact of mutations at these oversampled sites by plotting the change in log *K*_*D*_ (Δ log *K*_*D*_) at each site relative to the wild type. The red dashed line represents the global average Δ log *K*_*D*_ across the entire RBD.

In Figures 4e–j, each of these sites is selected either for its ability to escape one or more antibodies while maintaining ACE2 binding, as highlighted in Figure 4e. In particular, despite being trained on only a few percent of single-variant data, our model successfully identified critical antibody escape mutations that later emerged in major SARS-CoV-2 variants. These key positions include residue 484 (enabling LY-CoV016 and LY-CoV555 escape) in Omicron BA.1; residue 493 (enabling CoV016 and CoV555 escape) in Omicron BA.1, BA.2, and BA.5; positions 340 (enabling S309 escape) and 356 (enabling S309 escape) in Omicron BA.2; residue 445 (enabling REGN10987 escape) in variants B.1, BF.8, and XD; and position 486 (enabling CoV016 and CoV555 escape) in numerous variants ranging from N.6 to Omicron BA.5 to BL1. A comprehensive list of these positions and their associated variants can be found in the Supplementary Information. We also identified position 507 as an oversampled site. Although mutations at this position allow escape from the LY-CoV016 and REGN10987 antibodies, they significantly compromise the binding affinity of ACE2. This trade-off between immune escape and receptor binding likely explains why mutations at position 507 have not been widely observed in naturally circulating variants.

## Discussion

In this study, we developed an integrated framework VIRAL for identifying high-fitness SARS-CoV-2 RBD variants and predicting mutation-prone sites using minimal experimental data. Our approach combines three key components: pLM for sequence representation, Gaussian processes for efficient learning and uncertainty quantification, and biophysical modeling for fitness prediction. This integration demonstrates several significant advantages in the context of viral surveillance and variant prediction.

First, our model achieves high efficiency in identifying dangerous variants, obtaining up to 5-fold enrichment over random sampling while requiring experimental characterization of less than 1% of possible variants in the combinatorial mutagenesis landscape. This significant reduction in the number of required experiments could substantially accelerate the identification of concerning variants during early pandemic stages, when experimental resources are often limited. While initializing a model with some training data (as in this study) is advantageous, it is also feasible to start with zero training data, where zero-shot predictions initially carry equal uncertainty. Indeed, pLMs have been proved to be effective at zero-shot protein functions, provided they are trained on large and diverse protein sequence databases Meier et al. (2021). As more data are gathered, a sample-efficient model leveraging uncertainty can iteratively improve its predictions and confidence. This iterative cycle of computation and experimentation has been central to experiment prioritization, especially in drug discovery Eisenstein (2020).

Second, our comparative analysis of acquisition strategies reveals an important balance between exploitation and exploration in viral surveillance. While the greedy strategy efficiently samples known high-fitness regions, the UCB approach enables broader exploration of the sequence landscape, particularly into regions with higher semantic change. This broader sampling is crucial for identifying evolutionarily distant but potentially dangerous variants. The UCB strategy’s ability to discover multiple distinct clusters of high-fitness variants, as visualized in our UMAP analysis, demonstrates its value in maintaining diversity during the search process. This aligns with our understanding of viral evolution, where new variants often emerge by exploring antigenically novel regions while maintaining essential functionality such as binding and folding. Importantly, it would not be possible to reliably identify immune escape mutations based solely on their semantic change, as underlined in a recent study Allman et al. (2024). However, when combined with protein stability metrics and active learning frameworks, semantic change can play a complementary role in uncovering novel regions of the sequence landscape.

Here, we systematically sample diverse regions of the mutation space, uncovering high-risk variants with immune escape potential. Similarly, Luksza et al.’s predictive fitness model for influenza Łuksza and Lässig (2014) highlights how clades evolve by balancing antigenic novelty and fitness, ensuring escape from host immunity while maintaining functional integrity. Also, Meijer’s work Meijers et al. (2023) shows how population immunity, shaped by vaccinations and prior infections, drives the virus toward new antigenic regions, enabling it to escape cross-immunity and dominate through successive clade shifts. Together, these insights emphasize the importance of exploring antigenically distant regions to anticipate and mitigate emergent viral threats effectively.

Uncertainty sampling is just one of many active learning strategies that could be explored further. Other approaches, such as diversity-based sampling and expected predictive improvement, could enhance the robustness of the model. Hybrid strategies, which dynamically switch between acquisition functions, have been proposed in prior work. For instance, the DUAL framework (Donmez et al., 2007) adaptively alternates between uncertainty and representative-based sampling to improve model predictive power. Interestingly, prioritizing uncertainty in acquisition strategies for high fitness variant discovery is not universally adopted; for instance, Hie et al. Hie et al. (2020) penalized uncertainty when trying to discover compounds with nanomolar affinity for diverse kinases.

Third, our framework demonstrates high accuracy in identifying biologically relevant mutation sites. By defining the genetic space in terms of the effects of single mutations, we achieve a predictive advantage of two years compared to the baseline strategy, which relies on waiting for variants to emerge in nature, measuring their fitness, and then predicting the fitness effects of new mutations. Combining biophysics and active learning trained on *K*_*d*_ values offers two major advantages. First, *K*_*d*_ values can be experimentally measured early in a pandemic, unlike fitness values, which require the spread of mutations in the population to infer growth curves. Second, our biophysical fitness predictions are interpretable, unlike black-box models that directly output fitness values. Specifically, our predictions are driven by biophysical insights, such as antibody escape potential or tight ACE2 binding, making them iologically meaningful. Furthermore, the systematic oversampling of positions that persisted in variants of concern, particularly at sites 356, 484, 486 and 493, demonstrates that our model could highlight evolutionarily important sites using limited data.

A key assumption of our work is that the biophysical model serves as a reliable estimator of viral fitness, as demonstrated by Wang et al. Wang et al. (2024). Without the biophysical mapping from *K*_*D*_ to fitness space, active learning strategy could lead to the enrichment of variants that do not align with the correct fitness landscape. Fortunately, prior work by Cheron et al. and Rotem et al. Chéron et al. (2016); Rotem et al. (2018) demonstrated that fitness in RNA viruses can be quantitatively linked to molecular properties. These successes underscore the potential of biophysical approaches in modeling complex fitness landscapes for viruses beyond SARS-CoV-2, suggesting that our methodology could be generalized to address future pandemic threats.

## Methods

### Datasets

We utilize two distinct datasets in our research. The first is the combinatorial *K*_*D*_ measurements from the work of Moulana et al.Moulana et al. (2022, 2023) In their study, they systematically examined the interactions between all possible combinations of 15 mutations in the RBD of BA.1 relative to the Wuhan Hu-1 strain(totaling 32,768 genotypes) and ACE2, as well as four monoclonal antibodies (LY-CoV016, LY-CoV555, REGN10987, and S309).

The second dataset that we examined is a DMS from Starr et al. Starr et al. (2020, 2021a), providing for all possible RBD single mutants *K*_*d*_ values (ACE2) and escape ratios *ϵ*(*mut*) (LY-CoV016, LY-CoV555, REGN10987 and S309), and filtered on residues between 334 and 526 included. 28 sites are excluded from this dataset due to missing data for one of the biophysical constants. When computing AUC for comparison with GISAID data, we removed these sites from labels as well to ensure unbiased estimation.

Noting that the dissociation constant of RBD writes:

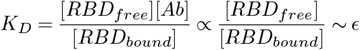

we assumed log dissociation constants of single mutants could be obtained as the sum of wild type log dissociation constant and the variation of escape ratio compared to its minimum value (wildtype):

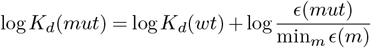

### ESM

ESM models are Transformer-based protein language models(pLM) designed to extract meaningful representations from protein sequences, enabling tasks such as structure prediction and large-scale protein characterization.

We use these models to obtain semantic representation of protein sequences. For a protein sequence of length *L*, we first describe it as a sequence of tokens 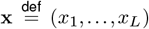. In the base of RBD, *L* = 201. We then run a forward pass of ESM and obtain the hidden representations of the final layer, (**h**_1_, …, **h**_*L*_), where each **h**_*i*_ ∈ ℝ^*K*^. Then, we use mean pooling of these vectors to obtain a representation of the entire sequence, 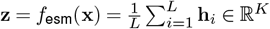. Finally, we re normalized the embeddings by mean and variance. We benchmarked different models: ESM1v Meier et al. (2021) (K=1280), ESM2 Lin et al. (2023) (K=1280), ESM3 Hayes et al. (2024) sequence only, and ESM3 sequence with structure information from PDB 6XF5 Zhou et al. (2020) (K=1536). Main results are obtained using ESM3 with structure encoding.

### Semantic Change

We can denote the sequence of wild-type RBD as **x**_*wt*_ and the mutant as **x**_*mt*_ where **x**_*mt*_ may have one or more different tokens than **x**_*wt*_. Semantic change is defined as the *L*_2_ norm of the embedding distance:

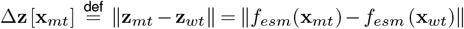

A high semantic change represents large change of the semantic meaning of the protein sequence, which we noticed to be correlated with decreased antibody binding affinity.

### Fitness and Biophysical Model

We determine the fitness of the RBD based on its contribution to viral infectivity, utilizing a biophysical model established by our previous work Wang et al. (2024). This model leverages the Boltzmann distribution to map molecular phenotypes—characterized by binding energies to cell receptors and antibodies—onto a fitness landscape. The RBD’s fitness is primarily determined by two factors: its binding affinity for the ACE2 receptor and its ability to evade antibody neutralization.

In our model, the RBD can exist in multiple states, each with its associated free energy: Unfolded state (*G*_*u*_), folded but unbound state (*G*_*f*_), folded and bound to ACE2 (*G*_*bA*_), folded and bound to one of four distinct antibodies (*G*_*ai*_, where *i* indexes the antibodies).

The fitness function can be expressed as:

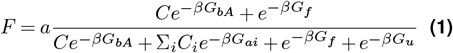

In this equation, 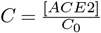 and 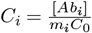 represent the normalized concentrations of ACE2 and antibodies respectively, where […] denotes molar concentration and *m*_*i*_ is a neutralization coefficient specific to each antibody. Noting that *β*Δ*G* = *β*(*G*_*i*_ − *G*_*f*_) = *ln*(*K*_*D*_)*/T* for every state *i*, where T is a hyperparameter proportional to system temperature, fitness can be expressed as a function of measured dissociation constants.

The model parameters—including the scaling factor *a* and effective concentrations *C* and *C*_*i*_—were calibrated by combining experimental measurements of dissociation constants (*K*_*D*_) with variant prevalence data from the GISAID database Elbe and Buckland-Merrett (2017a). This biophysical framework can then predict the fitness (*F*) of any RBD variant given its binding affinities, whether measured experimentally or predicted computationally.

In this paper we did not refit the biophysical model iteratively, instead we used the following coefficients fitted from et al.Wang et al. (2024): *T* = 1.6, *a* = 1.57363338, *C* = 5.4764 × 10^−7^, *K*_1_ = 5.6015 ×10^−8^, *K*_2_ = 4.5128 × 10^−8^, *K*_3_ = 7.1825 × 10^−8^, *K*_4_ = 4.7273 × 10^−7^. While swe did not refit these parameters iteratively, the model can be trained with very limited fitness data due to its low number of parameters and still approximate well fitness derived from population data.

### Uncertainty of predictions

To propagate variance from the predicted dissociation constants to the fitness function, we used standard error propagation formula, which states that the variance of a function *f* can be approximated as:

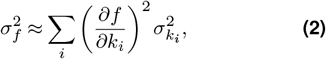

where 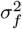 is the variance of the function, 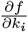 is the partial derivative of the function with respect to the *i*-th variable, and 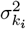 is the variance of the *i*-th variable. The variance of variables is obtained from the posterior distribution of the GP.

The derivatives 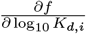 were computed symbolically using SymPy and evaluated numerically with the respective parameter values.

### Gaussian Process Kernel Selection

To model protein binding using Gaussian processes, we defined a kernel function that captures the notion of similarity between different variants in the sequence embedding space. We employed a Rational Quadratic kernel, which is well-suited for modeling functions with varying degrees of smoothness. The kernel function is defined as:

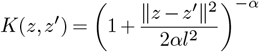

where *α* is the scale mixture parameter, and *l* is the length scale parameter.

From a Bayesian perspective, the kernel defines the prior covariance between data points, ensuring that variants that are closer in embedding space (∥*z* −*z*^′^∥^2^ small) exhibit similar binding values, while more distant variants remain weakly correlated. This reflects the fact that mutations with similar physicochemical properties and structural contexts tend to have correlated effects on protein binding.

In our framework, the kernel hyperparameters *α* and *l* were optimized by maximizing the marginal likelihood of the observed binding data, allowing the model to adaptively learn the appropriate scale of binding variation across the mutational landscape.

When training the Gaussian process on a training set (*Z*_1_, *Y*_1_) of size *N* and making predictions for a new point (*z*_2_, *y*_2_), we used the analytical solutions for the posterior distribution:

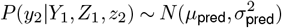

with

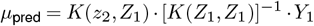

and

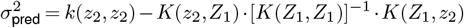

where *K*(*z*_2_, *Z*_1_) = *K*(*Z*_1_, *z*_2_)^*T*^ is a vector of dimension *N, K*(*Z*_1_, *Z*_1_) is the covariance matrix of dimension *N × N*, and *k*(*z*_2_, *z*_2_) is the kernel function evaluated at the test point.

The mean of the posterior distribution serves as a prediction for the output variable *y*_2_ corresponding to the input sample *z*_2_, while the variance (the diagonal of the covariance matrix) acts as a proxy for uncertainty. The mean of the posterior predictions in a Gaussian process represents a weighted average of the observed variables, with the weights determined by the covariance function.

In our specific application, *z* represents the embedding of a sequence, and *y*(*z*) = Δ log *K*_*D*_ denotes the variation of log-transformed dissociation constant compared to wildtype.

The kernel hyperparameters were optimized by maximizing the marginalized likelihood function using *sklearn*.*gaussianprocess* library.

### Model Benchmark

We evaluate the ability of VIRAL to rank variants by computing the Spearman correlation between inferred fitness scores and ground-truth labels. For the combinatorial dataset, the model is trained on varying numbers of data points, ranging from 20 to 160, and then used to predict the fitness of the remaining variants.

For the DMS dataset, training sets are constructed by sampling an average of *n* ∈ {0.2, 0.4, 0.6, 0.8, 1, 2, 3} mutations per site. When *n <* 1, a single mutant at each site is randomly selected and included in the training set with probability n.

To ensure robustness, each experiment is repeated 10 times for both datasets and for each training size, enabling a reliable assessment of the model’s predictive accuracy across different levels of data availability.

### Active Learning

The Bayesian optimization approach used in this study incorporates active learning principles to efficiently explore a discrete set of candidate sequences, referred to as the “pool.” This strategy selects and evaluates sequences in batches, progressively refining the optimization process. The iterative procedure consists of several key steps:

1. A random batch, denoted as *S*_0_, is initially selected from the candidate set *D*. The labels for these points is calculated (e.g., experimental dissociation constants), forming the initial training dataset *D*_*train*_. Unexplored dataset is then *D*^′^ ← *D*\*S*_0_
2. A surrogate model 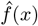 is then trained on the dataset *D*_*train*_, in order to predict fitness of unknown points in *D*^′^. Here, it first predicts dissociation constants, the latest being fed to a biophysical model which convert them into a fitness proxy. Each fitness prediction has value 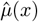 and variance 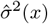.
3. An acquisition function 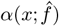 is used to determine the utility of acquiring a given point *x*. This function considers various metrics, such as the predicted values for fitness, as well as the associated uncertainties.
4. The points with the highest utility, denoted as an ensemble *X*^*^, are selected or “acquired”. We obtain the ground truth value for their dissociation constants and add them to the training set (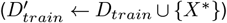 while removing it from the unexplored dataset (*D*^′^ ← *D*^′^\{*X*^*^}).

Steps 2-4 are repeated iteratively until a stopping criterion is met, such as a fixed number of iterations or insufficient improvement.

In our study, we adopt following active learning parameters. For the DMS benchmark we chose initial training size of 174 mutations (randomly sampled, one mutation per RBD site) corresponding to ∼5% of dataset and 10 more rounds acquiring 50 points each. For the Combinatorial benchmark, we chose initial training size of 20 variants (randomly sampled among single and double mutants) corresponding to ∼0.06% of dataset and 10 more rounds acquiring 10 points each. Active learning acquisition was repeated 10 times for each benchmark, using different initial training sets sampled as described.

### Acquisition Functions

Various acquisition functions for active learning are considered in this study, each influencing the point selection process differently. These include:

- Random 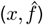: Randomly selecting points from the candidate set.
- Greedy 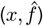: Selecting points based on the surrogate model’s mean 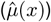.
- UCB(*x*): Employing the Upper Confidence Bound strategy, combining the mean and a scaled standard deviation 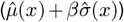.

To normalize the contribution of the variance term relative to the fitness values in UCB, we chose the coefficient *β* as:

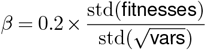

This ensures that the uncertainty term 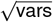 is scaled to have a comparable range to the fitness values, adjusted by a scaling. We explored in SI Figure S2 impact of uncertainty weight in UCB exploration.

#### Evaluation of Active Learning Performance

The primary metric used to evaluate our pipeline is the enrichment factor (EF), defined as

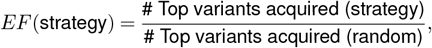

where “top variants” refers to those with fitness in the top *p*% of the dataset. Unless otherwise specified, we use p=10% as the default threshold for defining high-fitness variants. An EF value greater than 1 indicates that the strategy outperforms random sampling in identifying top variants.

Additionally, we compute the Area Under the Curve (AUC) using the active learning predicted fitness as a score to identify whether a variant belongs to the top 10% of the dataset. Notably, this metric is evaluated on the entire dataset (including both tested and untested variants) to ensure the results are not negatively biased against models that have acquired all the top variants. Lastly, we calculate the embedding variance of the tested variants as a quantitative measure of sequence exploration diversity. This metric is defined as the mean variance across all dimensions of the ESM3 embeddings among the acquired variants, providing intuition into how broadly the acquisition strategy samples the protein sequence space.

#### Comparison with GISAID data

We analyzed SARS-CoV-2 spike sequences from the GISAID database Elbe and Buckland-Merrett (2017b), collecting 15,371,428 sequences up to April 14, 2023. Following the methodology of Starr et al. Starr et al. (2021b), we implemented a sequence filtration process. Sequences were excluded if they were: (1) from non-human hosts, (2) outside the length range of 1,260-1,276 amino acids, (3) contained unicode errors, gaps, or ambiguous characters. The remaining sequences were aligned using MAFFT Katoh and Standley (2013).

The final dataset comprised 11,976,984 submissions, containing 25,725 unique RBD sequences. For each unique RBD sequence, we tracked its frequency of occurrence and estimated its emergence time using the 5th percentile of its temporal distribution. To ensure robustness, we excluded singleton sequences that appeared only once in the dataset.

We assessed site mutability by analyzing amino acid diversity at each position within the RBD. Sites were labeled as “highly mutable” if they exhibited at least 9 distinct amino acid variants out of the possible 20, observed in mutants with a total count of 10 or more. Using this cutoff of 10 on variant prevalence helps ensure that the identified mutations provided a clear evolutionary advantage rather than occurring randomly, allowing us to identify sites under selection pressure during the pandemic. To evaluate the effectiveness of our active learning pipeline, we used site sampling frequency as a performance metric and calculated the Area Under the Receiver Operating Characteristic Curve (AUC). The AUC score quantifies our algorithm’s ability to identify positions that emerged as mutation hotspots throughout the pandemic. AUC scores were also tested for mutation thresholds different from 9, see SI.

#### Baseline for identification of highly mutable sites

We define a baseline model to identify high-fitness mutations, trained on pandemic data. The training data includes mutations observed in the GISAID database within a variant having a minimum count of 1000, under the assumption that variants with counts below this threshold lack reliable fitness estimates. Specifically, mutations in variants with occurrences prior December 2020 / December 2022 and counts exceeding 1000 were selected. The baseline was trained on 14 mutations for 2021 deadline and 64 mutations for 2022 deadline. Single-batch acquisition size is 674 − training size, ensuring the total number of variants acquired by the active learning process and the baseline is the same. We then computed acquisition score for every site, based on this single batch acquisition, and compared it to GISAID data.

## Code Availability

The code used in this study will be made publicly available at https://github.com/m-huot/VIRAL upon manuscript acceptance. For access requests prior to publication, please contact the corresponding authors.

## Acknowledgments and Funding

This work is supported by NIH R35GM139571. The content is solely the responsibility of the authors and does not necessarily represent the official views of the National Institutes of Health. We gratefully acknowledge all data contributors, i.e., the Authors and their originating laboratories responsible for obtaining the specimens, and their submitting laboratories for generating the genetic sequence and metadata and sharing via the GISAID Initiative, on which this research is based.

## Notes

### Competing Interest Statement

The authors have declared no competing interest.

## Bibliography

Addetia, A., Piccoli, L., Case, J. B., Park, Y.-J., Beltramello, M., Guarino, B., Dang, H., de Melo, G. D., Pinto, D., Sprouse, K., et al. (2023). Neutralization, effector function and immune imprinting of omicron variants. Nature, 621(7979):592–601.

Allman, B., Vieira, L., Diaz, D. J., and Wilke, C. O. A systematic evaluation of the languageof-viral-escape model using multiple machine learning frameworks, (2024). URL http://biorxiv.org/lookup/doi/10.1101/2024.09.04.611278.

Barton, M. I., MacGowan, S. A., Kutuzov, M. A., Dushek, O., Barton, G. J., and Van Der Merwe, P. A. (2021). Effects of common mutations in the SARS-CoV-2 Spike RBD and its ligand, the human ACE2 receptor on binding affinity and kinetics. eLife, 10:e70658. doi: 10.7554/eLife.70658.

Carabelli, A. M., Peacock, T. P., Thorne, L. G., Harvey, W. T., Hughes, J., COVID-19 Genomics UK Consortium, De Silva, T. I., Peacock, S. J., Barclay, W. S., De Silva, T. I., Towers, G. J., and Robertson, D. L. (2023). SARS-CoV-2 variant biology: immune escape, transmission and fitness. Nature Reviews Microbiology. doi: 10.1038/s41579-022-00841-7.

Chéron, N., Serohijos, A. W. R., Choi, J.-M., and Shakhnovich, E. I. (2016). Evolutionary dynamics of viral escape under antibodies stress: A biophysical model. Protein Sci., 25 (7):1332–1340.

Donmez, P., Carbonell, J. G., and Bennett, P. N. Dual strategy active learning. In Kok, J.N., Koronacki, J., Mantaras, R.L.d., Matwin, S., Mladenič, D., and Skowron, A., editors, Machine Learning: ECML 2007, pages 116–127, Berlin, Heidelberg, (2007). Springer Berlin Heidelberg. ISBN 978-3-540-74958-5.

Eisenstein, M. (2020). Active machine learning helps drug hunters tackle biology. Nature Biotechnology, 38(5):512–514. doi: 10.1038/s41587-020-0521-4.

Elbe, S. and Buckland-Merrett, G. (2017). Data, disease and diplomacy: Gisaid’s innovative contribution to global health. Global challenges, 1(1):33–46.

Elbe, S. and Buckland-Merrett, G. (2017). Data, disease and diplomacy: GISAID’s innovative contribution to global health: Data, Disease and Diplomacy. Global Challenges, 1(1):33–46. doi: 10.1002/gch2.1018.

Gessner, A., Ober, S. W., Vickery, O., Oglić, D., and Uçar, T. Active learning for affinity prediction of antibodies, (2024). URL https://arxiv.org/abs/2406.07263.

Graff, D. E., Shakhnovich, E. I., and Coley, C. W. (2021). Accelerating high-throughput virtual screening through molecular pool-based active learning. Chem. Sci., 12:7866–7881. doi: 10.1039/D0SC06805E.

Hayes, T., Rao, R., Akin, H., Sofroniew, N. J., Oktay, D., Lin, Z., Verkuil, R., Tran, V. Q., Deaton, J., Wiggert, M., Badkundri, R., Shafkat, I., Gong, J., Derry, A., Molina, R. S., Thomas, N., Khan, Y., Mishra, C., Kim, C., Bartie, L. J., Nemeth, M., Hsu, P. D., Sercu, T., Candido, S., and Rives, A. Simulating 500 million years of evolution with a language model, (2024). URL http://biorxiv.org/lookup/doi/10.1101/2024.07.01.600583.

Hie, B., Bryson, B. D., and Berger, B. (2020). Leveraging Uncertainty in Machine Learning Accelerates Biological Discovery and Design. Cell Systems, 11(5):461–477.e9. doi: 10.1016/j.cels.2020.09.007.

Ito, J., Strange, A., Liu, W., Joas, G., Lytras, S., The Genotype to Phenotype Japan (G2P-Japan) Consortium, and Sato, K. A Protein Language Model for Exploring Viral Fitness Landscapes, (2024). URL http://biorxiv.org/lookup/doi/10.1101/2024.03.15.584819.

Katoh, K. and Standley, D. M. (2013). MAFFT Multiple Sequence Alignment Software Version 7: Improvements in Performance and Usability. Molecular Biology and Evolution, 30(4): 772–780. doi: 10.1093/molbev/mst010.

Khalak, Y., Tresadern, G., Hahn, D. F., De Groot, B. L., and Gapsys, V. (2022). Chemical Space Exploration with Active Learning and Alchemical Free Energies. Journal of Chemical Theory and Computation, 18(10):6259–6270. doi: 10.1021/acs.jctc.2c00752.

Lin, Z., Akin, H., Rao, R., Hie, B., Zhu, Z., Lu, W., Smetanin, N., Verkuil, R., Kabeli, O., Shmueli, Y., dos Santos Costa, A., Fazel-Zarandi, M., Sercu, T., Candido, S., and Rives, A. (2023). Evolutionary-scale prediction of atomic-level protein structure with a language model. Science, 379(6637):1123–1130. doi: 10.1126/science.ade2574.

Loux, T., Wang, D., and Shakhnovich, E. I. Does structural information improve esm3 for protein binding affinity prediction?

Maher, M. C., Bartha, I., Weaver, S., Di Iulio, J., Ferri, E., Soriaga, L., Lempp, F. A., Hie, B. L., Bryson, B., Berger, B., Robertson, D. L., Snell, G., Corti, D., Virgin, H. W., Kosakovsky Pond, S. L., and Telenti, A. (2022). Predicting the mutational drivers of future SARS-CoV-2 variants of concern. Science Translational Medicine, 14(633):eabk3445. doi: 10.1126/scitranslmed.abk3445.

Meier, J., Rao, R., Verkuil, R., Liu, J., Sercu, T., and Rives, A. Language models enable zero-shot prediction of the effects of mutations on protein function, (2021). URL http://biorxiv.org/lookup/doi/10.1101/2021.07.09.450648.

Meijers, M., Ruchnewitz, D., Eberhardt, J., łuksza, M., and Lässig, M. (2023). Population immunity predicts evolutionary trajectories of SARS-CoV-2. Cell, 186(23):5151–5164.e13. doi: 10.1016/j.cell.2023.09.022.

Moulana, A., Dupic, T., Phillips, A. M., Chang, J., Nieves, S., Roffler, A. A., Greaney, A. J., Starr, T. N., Bloom, J. D., and Desai, M. M. (2022). Compensatory epistasis maintains ACE2 affinity in SARS-CoV-2 omicron BA.1. Nat. Commun., 13(1):7011.

Moulana, A., Dupic, T., Phillips, A. M., Chang, J., Roffler, A. A., Greaney, A. J., Starr, T. N., Bloom, J. D., and Desai, M. M. (2023). The landscape of antibody binding affinity in SARS-CoV-2 omicron BA.1 evolution. Elife, 12.

Nabel, K. G., Clark, S. A., Shankar, S., Pan, J., Clark, L. E., Yang, P., Coscia, A., McKay, L. G. A., Varnum, H. H., Brusic, V., Tolan, N. V., Zhou, G., Desjardins, M., Turbett, S. E., Kanjilal, S., Sherman, A. C., Dighe, A., LaRocque, R. C., Ryan, E. T., Tylek, C., Cohen-Solal, J. F., Darcy, A. T., Tavella, D., Clabbers, A., Fan, Y., Griffiths, A., Correia, I. R., Seagal, J., Baden, L. R., Charles, R. C., and Abraham, J. (2022). Structural basis for continued antibody evasion by the SARS-CoV-2 receptor binding domain. Science, 375 (6578):eabl6251. doi: 10.1126/science.abl6251.

Obermeyer, F., Jankowiak, M., Barkas, N., Schaffner, S. F., Pyle, J. D., Yurkovetskiy, L., Bosso, M., Park, D. J., Babadi, M., MacInnis, B. L., et al. (2022). Analysis of 6.4 million sars-cov-2 genomes identifies mutations associated with fitness. Science, 376(6599):1327–1332.

Ozono, S., Zhang, Y., Ode, H., Sano, K., Tan, T. S., Imai, K., Miyoshi, K., Kishigami, S., Ueno, T., Iwatani, Y., Suzuki, T., and Tokunaga, K. (2021). SARS-CoV-2 D614G spike mutation increases entry efficiency with enhanced ACE2-binding affinity. Nature Communications, 12(1):848. doi: 10.1038/s41467-021-21118-2.

Romero, P. A., Krause, A., and Arnold, F. H. (2013). Navigating the protein fitness landscape with Gaussian processes. Proceedings of the National Academy of Sciences, 110(3). doi: 10.1073/pnas.1215251110.

Rotem, A., Serohijos, A. W. R., Chang, C. B., Wolfe, J. T., Fischer, A. E., Mehoke, T. S., Zhang, H., Tao, Y., Lloyd Ung, W., Choi, J.-M., Rodrigues, J. V., Kolawole, A. O., Koehler, S. A., Wu, S., Thielen, P. M., Cui, N., Demirev, P. A., Giacobbi, N. S., Julian, T. R., Schwab, K., Lin, J. S., Smith, T. J., Pipas, J. M., Wobus, C. E., Feldman, A. B., Weitz, D. A., and Shakhnovich, E. I. (2018). Evolution on the biophysical fitness landscape of an RNA virus. Mol. Biol. Evol., 35(10):2390–2400.

Seeger, M. (2004). Gaussian processes for machine learning. International journal of neural systems, 14(02):69–106.

Starr, T. N., Greaney, A. J., Hilton, S. K., Ellis, D., Crawford, K. H., Dingens, A. S., Navarro, M. J., Bowen, J. E., Tortorici, M. A., Walls, A. C., King, N. P., Veesler, D., and Bloom, J. D. (2020). Deep Mutational Scanning of SARS-CoV-2 Receptor Binding Domain Reveals Constraints on Folding and ACE2 Binding. Cell, 182(5):1295–1310.e20. doi: 10.1016/j.cell.2020.08.012.

Starr, T. N., Czudnochowski, N., Liu, Z., Zatta, F., Park, Y.-J., Addetia, A., Pinto, D., Beltramello, M., Hernandez, P., Greaney, A. J., Marzi, R., Glass, W. G., Zhang, I., Dingens, A. S., Bowen, J. E., Tortorici, M. A., Walls, A. C., Wojcechowskyj, J. A., De Marco, A., Rosen, L. E., Zhou, J., Montiel-Ruiz, M., Kaiser, H., Dillen, J. R., Tucker, H., Bassi, J., Silacci-Fregni, C., Housley, M. P., Di Iulio, J., Lombardo, G., Agostini, M., Sprugasci, N., Culap, K., Jaconi, S., Meury, M., Dellota Jr, E., Abdelnabi, R., Foo, S.-Y. C., Cameroni, E., Stumpf, S., Croll, T. I., Nix, J. C., Havenar-Daughton, C., Piccoli, L., Benigni, F., Neyts, J., Telenti, A., Lempp, F. A., Pizzuto, M. S., Chodera, J. D., Hebner, C. M., Virgin, H. W., Whelan, S. P. J., Veesler, D., Corti, D., Bloom, J. D., and Snell, G. (2021). SARS-CoV-2 RBD antibodies that maximize breadth and resistance to escape. Nature, 597(7874): 97–102. doi: 10.1038/s41586-021-03807-6.

Starr, T. N., Greaney, A. J., Addetia, A., Hannon, W. W., Choudhary, M. C., Dingens, A. S., Li, J. Z., and Bloom, J. D. (2021). Prospective mapping of viral mutations that escape antibodies used to treat COVID-19. Science, 371(6531):850–854. doi: 10.1126/science.abf9302.

Tuekprakhon, A., Nutalai, R., Dijokaite-Guraliuc, A., Zhou, D., Ginn, H. M., Selvaraj, M., Liu, C., Mentzer, A. J., Supasa, P., Duyvesteyn, H. M., Das, R., Skelly, D., Ritter, T. G., Amini, A., Bibi, S., Adele, S., Johnson, S. A., Constantinides, B., Webster, H., Temperton, N., Klenerman, P., Barnes, E., Dunachie, S. J., Crook, D., Pollard, A. J., Lambe, T., Goulder, P., Paterson, N. G., Williams, M. A., Hall, D. R., Fry, E. E., Huo, J., Mongkolsapaya, J., Ren, J., Stuart, D. I., Screaton, G. R., Conlon, C., Deeks, A., Frater, J., Frending, L., Gardiner, S., Jämsén, A., Jeffery, K., Malone, T., Phillips, E., Rothwell, L., and Stafford, L. (2022). Antibody escape of SARS-CoV-2 Omicron BA.4 and BA.5 from vaccine and BA.1 serum. Cell, 185(14):2422–2433.e13. doi: 10.1016/j.cell.2022.06.005.

Upadhyay, V., Panja, S., Lucas, A., Patrick, C., and Mallela, K. M. (2023). Biophysical evolution of the receptor-binding domains of sars-covs. Biophysical Journal, 122(23): 4489–4502.

Wang, D., Huot, M., Mohanty, V., and Shakhnovich, E. I. (2024). Biophysical principles predict fitness of sars-cov-2 variants. Proceedings of the National Academy of Sciences, 121(23):e2314518121. doi: 10.1073/pnas.2314518121.

Wang, Q., Iketani, S., Li, Z., Liu, L., Guo, Y., Huang, Y., Bowen, A. D., Liu, M., Wang, M., Yu, J., Valdez, R., Lauring, A. S., Sheng, Z., Wang, H. H., Gordon, A., Liu, L., and Ho, D. D. (2023). Alarming antibody evasion properties of rising SARS-CoV-2 BQ and XBB subvariants. Cell, 186(2):279–286.e8. doi: 10.1016/j.cell.2022.12.018.

Zhang, Q. E., Lindenberger, J., Parsons, R. J., Thakur, B., Parks, R., Park, C. S., Huang, X., Sammour, S., Janowska, K., Spence, T. N., et al. (2024). Sars-cov-2 omicron xbb lineage spike structures, conformations, antigenicity, and receptor recognition. Molecular Cell, 84(14):2747–2764.

Zhou, T., Teng, I.-T., Olia, A. S., Cerutti, G., Gorman, J., Nazzari, A., Shi, W., Tsybovsky, Y., Wang, L., Wang, S., Zhang, B., Zhang, Y., Katsamba, P. S., Petrova, Y., Banach, B. B., Fahad, A. S., Liu, L., Lopez Acevedo, S. N., Madan, B., Olivera De Souza, M., Pan, X., Wang, P., Wolfe, J. R., Yin, M., Ho, D. D., Phung, E., DiPiazza, A., Chang, L., Abiona, O., Corbett, K. S., DeKosky, B. J., Graham, B. S., Mascola, J. R., Misasi, J., Ruckwardt, T., Sullivan, N. J., and Shapiro, L. (2020). Structure-Based Design with Tag-Based Purification and In-Process Biotinylation Enable Streamlined Development of SARS-CoV-2 Spike Molecular Probes. SSRN Electronic Journal. doi: 10.2139/ssrn.3639618.

łuksza, M. and Lässig, M. (2014). A predictive fitness model for influenza. Nature, 507 (7490):57–61. doi: 10.1038/nature13087.

