## Supplementary Information for "Few-Shot Viral Variant Detection via Bayesian Active Learning and Biophysics"

### Supplementary Materials for:
### Few-Shot Viral Variant Detection via Bayesian Active Learning and Biophysics

Huot and Wang et al.

#### Supplementary information

This file includes:

Section S1. Active learning on DMS
Section S2. Influence of uncertainty weight in UCB acquisition
Section S3. Identification of mutation-prone sites

##### S1. Active learning on DMS

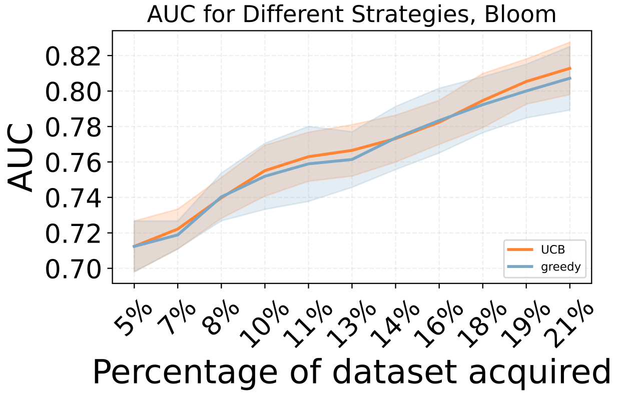


(a)

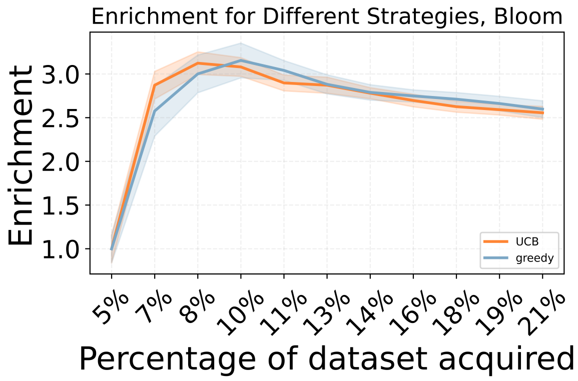


(b)

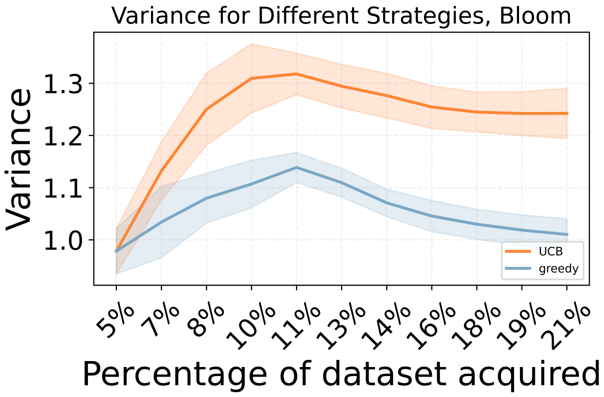


(c)

Fig. S.1. Active learning on DMS. (a) Area Under the Curve (AUC), representing the model's ability to identify top fitness variants, is shown for different strategies (UCB and greedy). An AUC above 0.5 indicates effective identification of high-fitness variants. Each round corresponds to acquiring a new batch of variants, improving the predictor. Shaded regions represent standard deviations across runs. (b) Enrichment in top variants across acquisition runs for each strategy. Values above 1 represent improvement other a random acquisition. (c) Comparison of embedding variance of points acquired across rounds between UCB (orange) and greedy (blue) strategies

We assess the capacity of our pipeline combined with active learning to identify high fitness single variants in the DMS dataset. This task requires to model the impact of approximately $193 * 19$ mutations, with initial knowledge of only one mutation per site. Interestingly, the UCB acquisition perform best in early rounds, while greedy acquisition gets better as the predictor gets more accurate on the longer term. This is explained by the high accuracy of our model to identify top variants after a few round of acquisitions, with AUC above 0.80.

#### S2. Influence of uncertainty weight in UCB acquisition

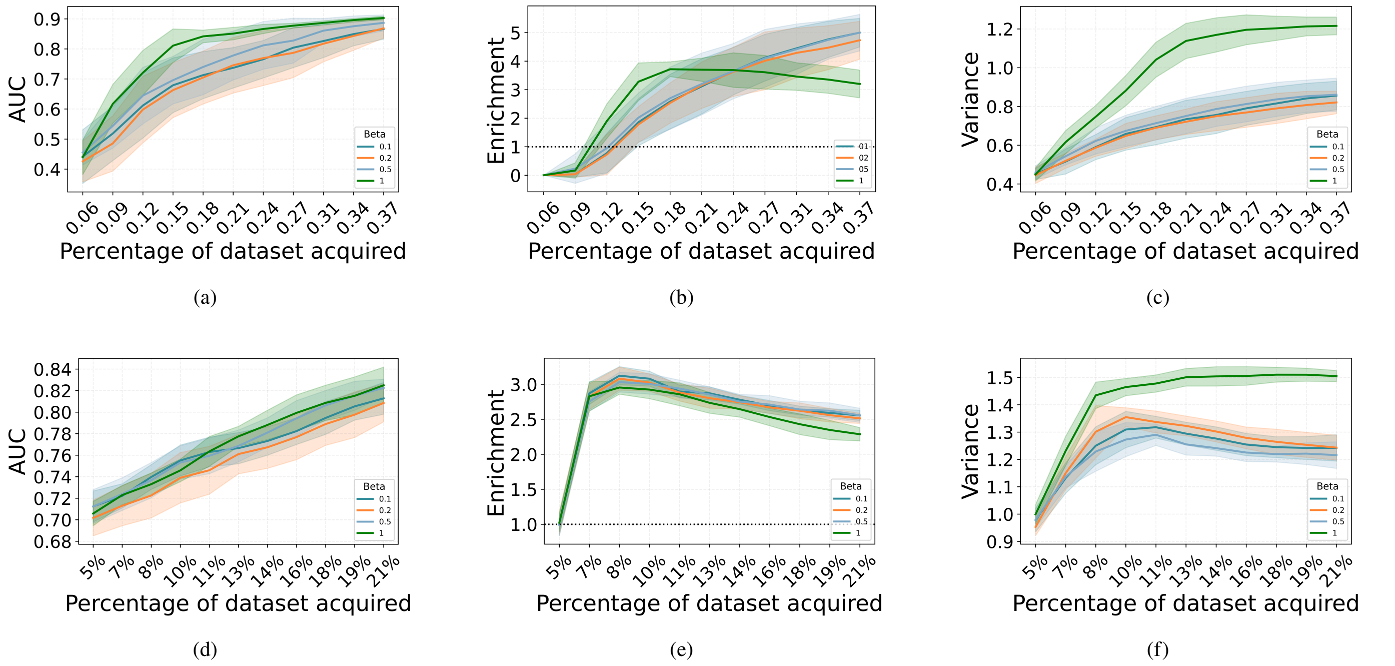


Fig. S.2. Influence of uncertainty weight $\beta$ in UCB acquisition. (a-c) AUC, enrichment, and embedding variance on the CM dataset. (d-f) Similar analysis on the DMS dataset. Different values of $\beta$ control the balance between exploration and exploitation, affecting model performance in identifying high-fitness variants.

Increasing the uncertainty weight $\beta$ in UCB acquisition enhances embedding variance and higher diversity in the training set and potentially improving predictive performance (higher AUC). However, in the long run, as the predictor becomes more accurate, it is preferable to reduce the emphasis on uncertainty, allowing the model to rely more on its predictions for optimal decision-making. Consequently, we selected $\beta = 0.2$ as a reasonable trade-off between exploitation and exploration of the mutational landscape. However, it is important to note that the impact of $\beta$ on overall performance remains relatively limited, and one can choose any reasonable $\beta$ *a priori* for the purpose of pandemic prevention.

$\beta$ is then scaled by $\frac{\text{std(fitnesses)}}{\text{std}(\sqrt{\text{vars}})}$ to ensure that the uncertainty term $\sqrt{\text{vars}}$ has a comparable range to the fitness values

#### *S3. Identification of mutation-prone sites*

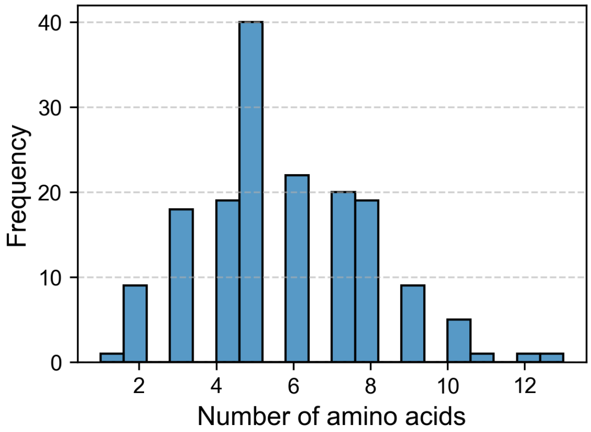


(a)

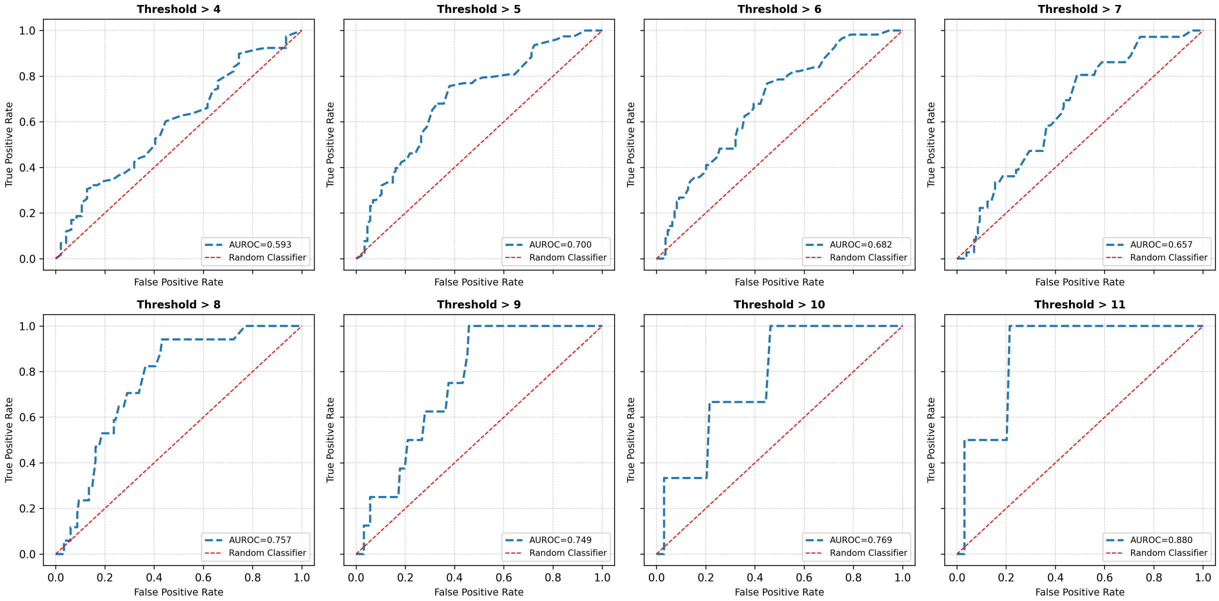


(b)

Fig. S.3. Active learning performance for identifying mutation-prone sites across different thresholds. (a) Histogram showing the distribution of SARS-CoV-2 RBD sites by the number of distinct amino acid substitutions observed during the pandemic. The data reveals most sites experienced between 4-8 different amino acid changes, with a peak at 6 amino acid variants. (b) ROC curves demonstrating VIRAL's ability to identify highly mutable sites in the GISAID database across various thresholds (from >4 to >11 mutations). The averaged acquisition scores from 10 independent runs show consistently strong predictive performance, with AUC values ranging from 0.593 to 0.890. Higher thresholds (>9, >10, >11) yield the best performance (AUC>0.75), indicating VIRAL is particularly effective at identifying the most highly mutable positions in the viral genome.

We assessed site mutability by analyzing amino acid diversity at each position within the RBD. In the main text, sites were labeled as "highly mutable" if they exhibited at least 9 distinct amino acid variants out of the possible 20, observed in mutants with a count of 10 or more. As demonstrated in Figure S3, the performance of VIRAL in identifying highly mutated sites remains robust across different thresholds defining mutation frequency (from >4 to >11 mutations), confirming the consistency of our approach regardless of the specific cutoff chosen for analysis.
